# Role of Complexin 2 in the regulation of hormone secretion from the islet of Langerhans

**DOI:** 10.1101/2024.10.28.620710

**Authors:** Xue Wen Ng, Michael R. DiGruccio, Chen Kong, Jeongmin Lee, David W. Piston

## Abstract

Regulated secretion of insulin from β-cells, glucagon from α-cells, and somatostatin from δ-cells is necessary for the maintenance of glucose homeostasis. The release of these hormones from pancreatic islet cells requires the assembly and disassembly of the SNARE protein complex to control vesicle fusion and exocytosis. Complexin 2 (Cplx 2) is a small soluble synaptic protein that participates in the priming and release steps of vesicle fusion. It plays a dual role as a molecular switch that first clamps and prevents fusion pore opening, and subsequently undergoes a conformational change upon Ca^2+^ binding to synaptotagmin to facilitate exocytosis. Using a Cplx 2 knockout (KO) mouse model, we show a direct inhibitory role of Cplx 2 for glucagon and somatostatin secretion, along with an indirect role in the paracrine inhibition of insulin secretion by somatostatin. Deletion of Cplx 2 increases glucagon and somatostatin secretion from intact mouse islets, while there is no difference in insulin secretion between WT and Cplx 2 KO islets. The normal paracrine inhibition of insulin secretion by somatostatin is disrupted in Cplx 2 KO islets. On the contrary, deletion of Cplx 2 did not affect the known role of somatostatin in the paracrine inhibition of glucagon at elevated glucose levels, since the paracrine inhibition of glucagon secretion by somatostatin is similar for both WT and Cplx 2 KO islets. In both β- and α-cells, the secretion profiles are parallel to Ca^2+^ activity changes following somatostatin treatment of WT and Cplx 2 KO islets. The loss of paracrine inhibition of insulin secretion is substantiated by direct measurements of insulin vesicle fusion events in Cplx 2 KO islets. Together, these data show a differential role for Cplx 2 in regulating hormone secretion from pancreatic islets.

## Introduction

Blood glucose homeostasis requires the proper regulation of secretion of insulin, glucagon and somatostatin from pancreatic islet β-, α- and δ-cells, respectively. The secretion mechanism of these islet secretory hormones and factors is driven by an intricate network of interconnected pathways. In each cell type, signaling begins with an influx of glucose via transporters (GLUTs), glucokinase-mediated metabolism of glucose through glycolysis and mitochondrial metabolism. Increased metabolic flux leads to an increase in ATP:ADP ratio, which triggers closure of ATP-sensitive K^+^ channels (K_ATP_) and membrane depolarization. While the details remain controversial, depolarization causes activation (β- and δ-cells) or perhaps deactivation (α-cells) of voltage-gated Ca^2+^ channels. Ca^2+^ influx and activity are required for exocytosis, and it stimulates insulin and somatostatin secretion. α-cell Ca^2+^ activity is heterogeneous, but any decrease in Ca^2+^ activity of these cells is expected to inhibit glucagon secretion (1, 2). Islet cell secretion is not only regulated by these intrinsic mechanisms, but also via intercellular communication within the islet. Cell-to-cell messages are mediated by paracrine signaling, where a factor released by neighboring cells interacts with receptors in the target cell to enhance or inhibit secretion, and juxtacrine signaling through direct contacts between plasma membrane linked ligands and receptors (1). Crucially, dysregulation of insulin and glucagon secretory pathways is observed in diabetes where there is diminished insulin secretion and elevated glucagon secretion (hyperglucagonemia) in response to glucose (3).

The release of islet hormones is controlled by molecular mechanisms at the plasma membrane similar to synaptic exocytosis. Soluble N-ethylmaleimide-sensitive factor attachment protein receptor (SNARE) proteins, Sec1/Munc18-like (SM) proteins and other associated proteins work collectively to mediate the assembly and disassembly of the four-helical SNARE protein complex bundle to facilitate synaptic vesicle docking, priming, fusion and release between the vesicle membrane and the plasma membrane (4). During exocytosis, secretory vesicles dock onto the plasma membrane and the SNAP/SNARE complexes on the vesicular and plasma membranes are primed for fusion. During the priming step, complexin (Cplx) binds to the prefusion trans-SNARE protein complex and acts as a molecular switch to clamp the trans-SNARE protein complex. Cplx undergoes a conformational change, evoked by calcium binding to Ca^2+^ sensor synaptotagmin, which releases the clamped trans-SNARE protein complex for binding with synaptotagmin, thus initiating fusion pore opening (5–7).

Complexins are a family of soluble small synaptic proteins that bind to the prefusion SNARE protein complex to facilitate vesicle fusion and exocytosis. There are four known isoforms of complexins (Cplxs 1 – 4) (8). Cplxs 1 and 2 are expressed in the brain and endocrine tissues, with significant expression in the pancreas, while Cplxs 3 and 4 are expressed mainly in the eye and brain (9). Cplxs have a crucial dual role in regulating Ca^2+^-dependent vesicle fusion by acting as a molecular switch that clamps and activates different stages of the SNARE protein complex assembly (5, 6). The central helix of Cplx first binds to the SNARE protein complex between the VAMP2 and syntaxin-1 helices (10). In the clamped state, the accessory helix of Cplx prevents fusion pore opening. Upon Ca^2+^ binding to synaptotagmin, Cplx undergoes a rearrangement where its accessory helix parallels the SNARE protein complex that allows synaptotagmin to bind to the SNARE complex, release the clamp, and activate fusion pore opening. Cplxs have been shown to exhibit both inhibitory (negative regulator) and stimulatory (positive regulator) effects on exocytosis *in vitro* and *in vivo*. Several studies have demonstrated that Cplxs act as negative regulators (clamp) of vesicle fusion (11–16). On the other hand, there are studies supporting the positive regulatory role (facilitator) of Cplxs in vesicle fusion (17–19). These contradictory findings can be explained by a recent study that establishes Cplx as both a checkpoint protein to ensure proper SNARE protein complex assembly during vesicle priming and a facilitator of vesicle fusion (20).

Cplx 1 was established as both a negative and positive regulator of glucose-stimulated insulin secretion in rat INS1 and mouse βTC3 cell lines, where both overexpression and small interfering RNA (siRNA) knockdown of Cplx 1 led to the decrease in glucose-stimulated insulin secretion (21). However, little is known on how Cplx 2 affects pancreatic islet cell hormone secretion. We examined the role of Cplx 2 in regulating glucose-induced insulin, glucagon, and somatostatin secretion. We demonstrate that both glucagon and somatostatin secretion are elevated in Cplx 2 knockout (KO) mouse islets as compared to wild-type (WT) islets, suggesting an inhibitory role of Cplx 2 on vesicle fusion in α- and δ-cells. This effect was not observed in β-cells where insulin secretion remains unchanged with the loss of Cplx 2 in intact mice islets. Upon administering somatostatin, however, we observe a significant loss of paracrine inhibition of β-cell insulin secretion from intact Cplx 2 KO mice islets. Conversely, paracrine inhibition of glucagon secretion in α-cells by somatostatin remained intact and similar for both Cplx 2 KO and WT islets. Loss of somatostatin paracrine inhibition of insulin secretion in β-cells of Cplx 2 KO mice islets is consistent with measurements of β-cell Ca^2+^ activity and insulin vesicle fusion events in these islets. Our findings suggest that Cplx 2 plays an important role as a negative regulator of vesicle fusion in pancreatic islet α- and δ-cells, and its clamping effect extends to the regulation of somatostatin paracrine inhibition of insulin secretion through β-cell Ca^2+^ activity in intact mouse islets.

## Results

### Complexin 2 is expressed in pancreatic β-, α- and δ-cells

*Cplx 1* RNA and protein expressions have been demonstrated in pancreatic β-cells (21), but the expression of Cplx 2 in the pancreatic islet remains unknown. We first validated the expression of *Cplx 2* RNA in β-, α- and δ-cells of fixed wild-type (WT) mouse islets by RNA fluorescence *in-situ* hybridization (RNA-FISH). RNA-FISH was performed in fixed WT mouse islets with the RNAScope kit and custom-designed *Mus musculus* Cplx 2 (Mm-Cplx 2) probe specific to the detection of *Cplx 2* RNA while avoiding other homologous genes. The Cplx 2 RNA-labeled fixed WT mice islets were co-immunostained with antibodies against insulin, glucagon or somatostatin to identify β-, α- or δ-cells, respectively. We observe *Cplx 2* RNA expression in pancreatic β-, α- and δ-cells (Fig. 1A) and quantify RNA copy/cell for all cell types (Fig. S1A). Cplx 2 protein expression was measured in fixed pancreatic islet sections of WT and Cplx 2 KO mice. Immunofluorescence imaging was performed on fixed pancreatic islet sections of WT and Cplx 2 KO mice with antibody against Cplx 2 co-immunostained with antibodies against either insulin, glucagon or somatostatin for the identification of β-, α- or δ-cells, respectively. Cplx 2 protein expression is observed in pancreatic β-, α- and δ-cells of WT pancreatic islet sections but not in pancreatic β-, α- and δ-cells of Cplx 2 KO pancreatic islet sections (Fig. 1B and Fig. S1B).

**Figure 1.**
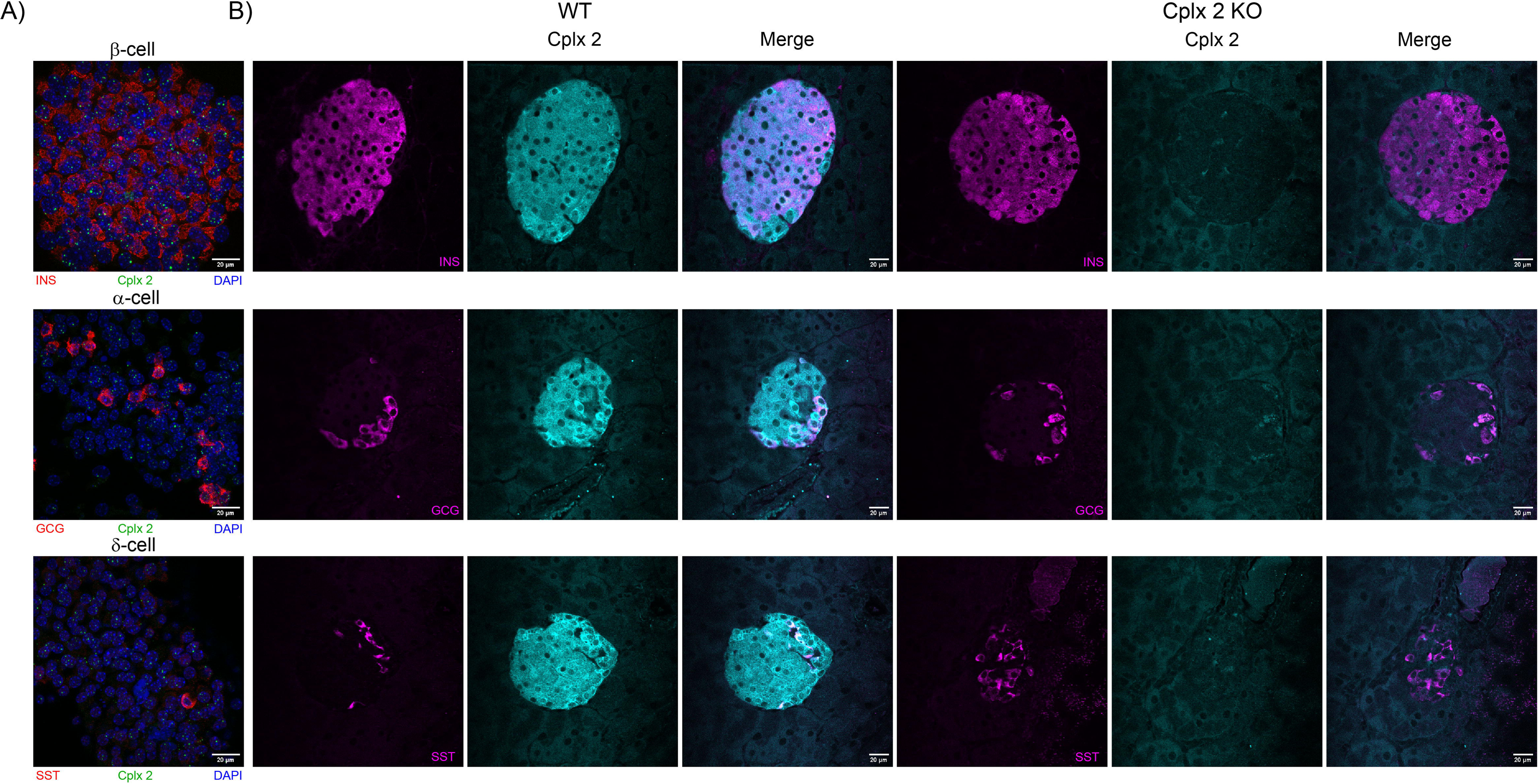
Cplx 2 is expressed in mouse pancreatic islet β-, α- and δ-cells, and loss of Cplx 2 protein expression is observed in Cplx 2 KO mouse islets. (A) Representative images of RNA fluorescence *in-situ* hybridization (FISH) of *Cplx 2* (green puncta) transcripts in fixed WT mouse islets co-immunostained with insulin (INS) for β-cells (upper panel), glucagon (GCG) for α-cells (middle panel), and somatostatin (SST) for δ-cells (bottom panel). (B) Comparison of Cplx 2 protein expression in fixed WT and Cplx 2 KO mouse pancreatic islet sections immunostained with antibodies against Cplx 2 (cyan) and either insulin (INS), glucagon (GCG) or somatostatin (SST) (magenta). Scale bars are 20 µm.

### Complexin 2 acts as a clamp for glucagon and somatostatin secretions in intact pancreatic islet

Cplx 1 was shown to both positively and negatively regulate glucose-dependent insulin secretion in pancreatic β-cells (21). Upon verifying the presence of Cplx 2 in pancreatic β-, α- and δ-cells, we investigated the role of Cplx 2 in regulating the secretions of all three islet cell types in both intact and dispersed mice islets. Insulin, glucagon and somatostatin secretion measurements are shown from both intact and dispersed WT and Cplx 2 KO mice islets at 1 mM (low) and 11 mM (high) glucose levels (Fig. 2). While insulin secretion remains unchanged between intact WT and KO islets (Fig. 2A), glucagon secretion at low glucose and somatostatin secretion at both low and high glucose are elevated in KO islets compared to WT islets (Fig. 2B and C). In intact islets, loss of Cplx 2 does not disrupt the glucose response of insulin, glucagon and somatostatin secretions (Fig. 2A – C), indicating that Cplx 2 has an auxiliary role in maintaining proper glucose homeostasis. We tested the effects of Cplx 2 KO in dispersed mouse islet cells which lack paracrine and juxtacrine interactions, and therefore have different glucose response in their secretion profiles than intact mouse islets (1). Similar to intact islets, insulin secretion in dispersed Cplx 2 KO islet cells is comparable to dispersed WT islet cells (Fig. 2D). It is worth noting that although the insulin secretion of dispersed Cplx 2 KO islet cells between low and high glucose levels are statistically less significant (P = 0.0562), its glucose response matches that of the dispersed WT islet cells (Fig. 2D). The clamping role of Cplx 2 on glucagon secretion is observed at low glucose level in dispersed cells (Fig. 2E). As expected, the glucose response of glucagon secretion is reversed in dispersed islet cells as compared to intact islets (Fig. 2B and E). The inhibitory role of Cplx 2 for somatostatin secretion is not seen for dispersed mouse islet cells (Fig. 2F), suggesting that Cplx 2 regulation in pancreatic islets requires intercellular communication. Taken together, these data establish an inhibitory role of Cplx 2 on glucagon and somatostatin secretions in intact mouse islets.

**Figure 2.**
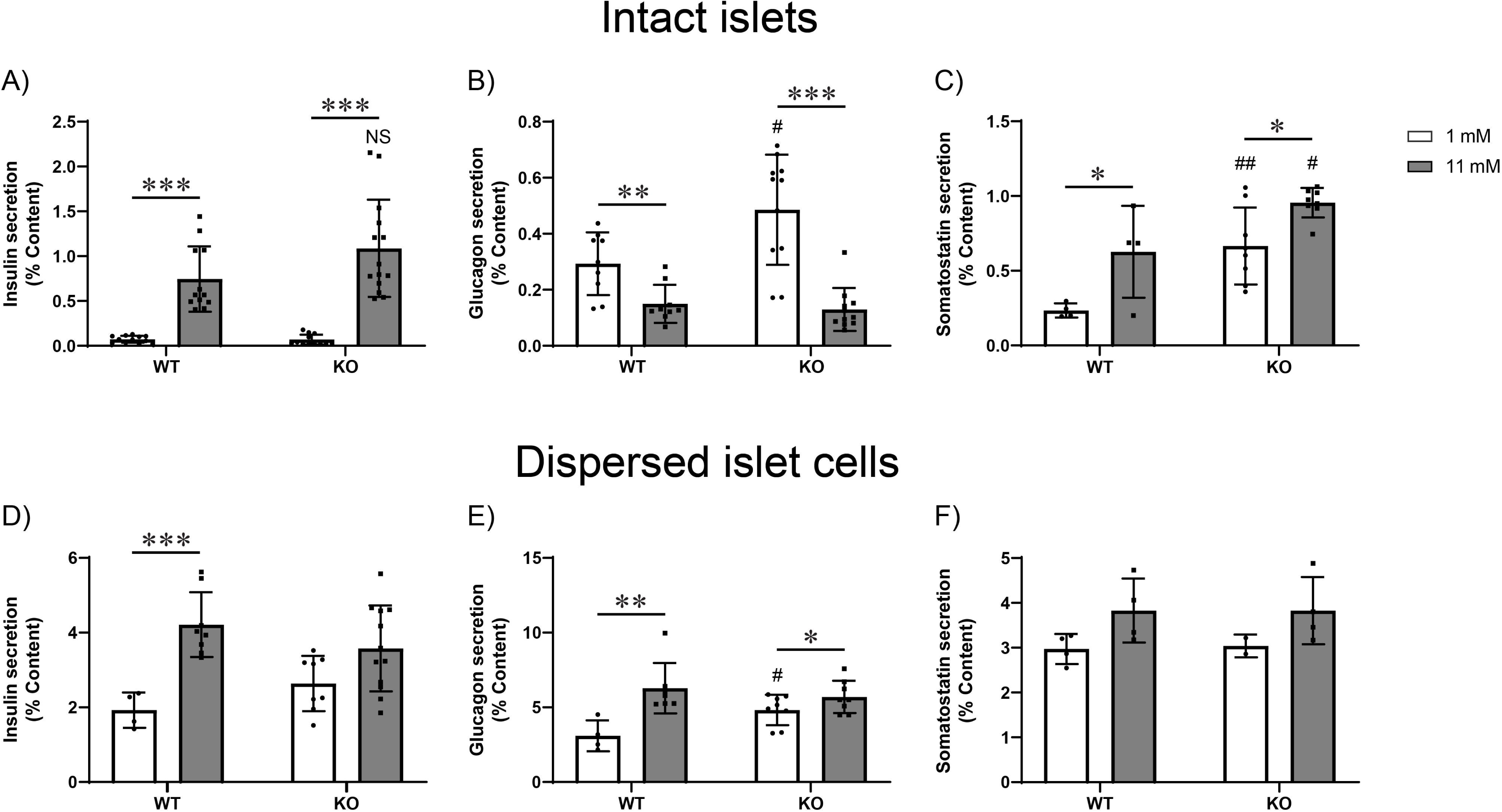
Effect of loss of Cplx 2 on insulin, glucagon and somatostatin secretion from intact mouse islets and dispersed islet cells. Insulin (A), glucagon (B) and somatostatin (C) secretions were measured in wild-type (WT) and Cplx 2 KO (KO) intact mouse islets at low (1 mM, white) and high (11 mM, gray) glucose concentrations. Insulin (D), glucagon (E) and somatostatin (F) secretions were measured in WT and KO dispersed cells at low (1 mM, white) and high (11 mM, gray) glucose concentrations. Data presented as mean ± SD (n > 2 experiments). Statistical significance between different glucose concentrations either in WT or KO islets was determined using unpaired *t* test ***P < 0.001; **P < 0.01; *P < 0.05. Statistical significance between WT and KO at the same glucose concentration was measured with unpaired *t* test ##P < 0.01; #P < 0.05.

### Loss of Complexin 2 leads to disrupted paracrine inhibition of insulin secretion by somatostatin in intact mouse islets

Upon ascertaining the clamping role of Cplx 2 in intact mouse islets, we investigated the effects of the loss of Cplx 2 on intact islets and dispersed islet cells treated with somatostatin (SST). SST activates receptors (SSTRs) on β- and α-cells, which inhibits insulin and glucagon secretion (1, 22). Intact WT and Cplx 2 KO mouse islets were treated with 1 μM SST at both low (1 mM) and high (11 mM) glucose concentrations and their insulin and glucagon secretions were measured. Treatment with SST on WT islets leads to significant inhibition of insulin release at 11 mM glucose concentration (Fig. 3A). On the contrary, SST treatment did not inhibit glucose-dependent insulin secretion from Cplx 2 KO islets at either glucose level (Fig. 3A). These data indicate that the inhibitory role of Cplx 2 associates with SST signaling at high glucose levels. Glucagon secretion was also measured in both WT and Cplx 2 KO mouse islets at 1 mM and 11 mM glucose levels in the absence and presence of 1 μM SST. Treatment with 1 μM SST on WT islets suppresses glucagon secretion at low glucose (Fig. 3B). In contrast to insulin secretion following SST-treatment, we do not see impairment of paracrine inhibition of glucagon secretion by SST in Cplx 2 KO islets (Fig. 3B).

**Figure 3.**
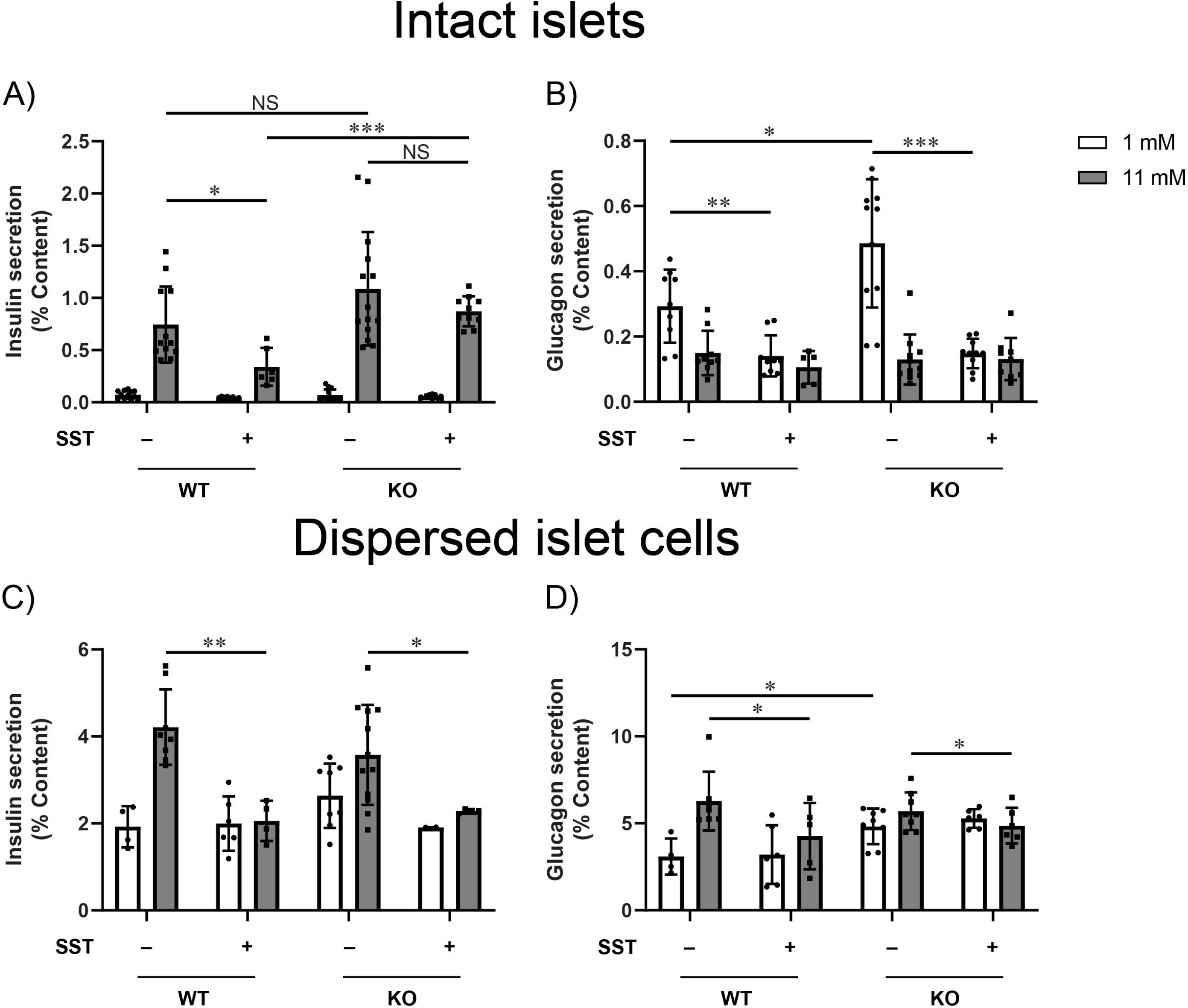
Loss of Cplx 2 leads to deficiency in paracrine inhibition of insulin secretion by somatostatin. Insulin (A) and glucagon (B) secretion from WT and Cplx 2 KO (KO) intact mouse islets at low (1 mM, white) and high (11 mM, gray) glucose concentrations with or without 1 µM somatostatin (SST) treatment. Insulin (C) and glucagon (D) secretion from WT and KO dispersed cells at low (1 mM, white) and high (11 mM, gray) glucose concentrations with or without 1 µM SST treatment. Data presented as mean ± SD (n > 2 experiments). Statistical significance between conditions was determined using unpaired *t* test ***P < 0.001; **P < 0.01; *P < 0.05.

We subsequently performed secretion experiments on dispersed WT and Cplx 2 KO islet cells treated with 1 μM SST. Both dispersed WT and Cplx 2 KO islet cells demonstrate inhibition of insulin (Fig. 3C) and glucagon (Fig. 3D) secretion by SST treatment at high glucose levels. These data suggest that loss of Cplx 2 has little effect on the secretion from islet cells that lack paracrine and juxtacine signaling mechanisms.

### Inhibitory role of Cplx 2 occurs downstream of paracrine receptor activation in intact mouse islets

Since it was demonstrated that loss of Cplx 2 disrupts the paracrine inhibition of insulin secretion by SST, we sought to determine whether such disruption occurs by antagonizing membrane receptors that drive paracrine signaling. Insulin and glucagon secretion was quantified from WT and Cplx 2 KO islets at low (1 mM) and high (11 mM) glucose levels with or without 2 μM S961, an insulin receptor (IR) antagonist, or 100 nM CYN 154806, an SSTR2 antagonist. We observe no difference in the response of insulin and glucagon release to IR or SSTR2 antagonism for intact WT and Cplx 2 KO islets (Fig. 4A and B). This indicates that Cplx 2 acts downstream from membrane receptor activation. Similar insulin and glucagon secretion experiments were conducted for dispersed WT and Cplx 2 KO mouse islet cells at 1 mM and 11 mM glucose levels with or without 2 μM S961 or 100 nM CYN 154806 treatment, and the outcomes were comparable to those in intact islets, emphasizing the lack of Cplx 2 action at the membrane receptor activation step (Fig. 4 C and D).

**Figure 4.**
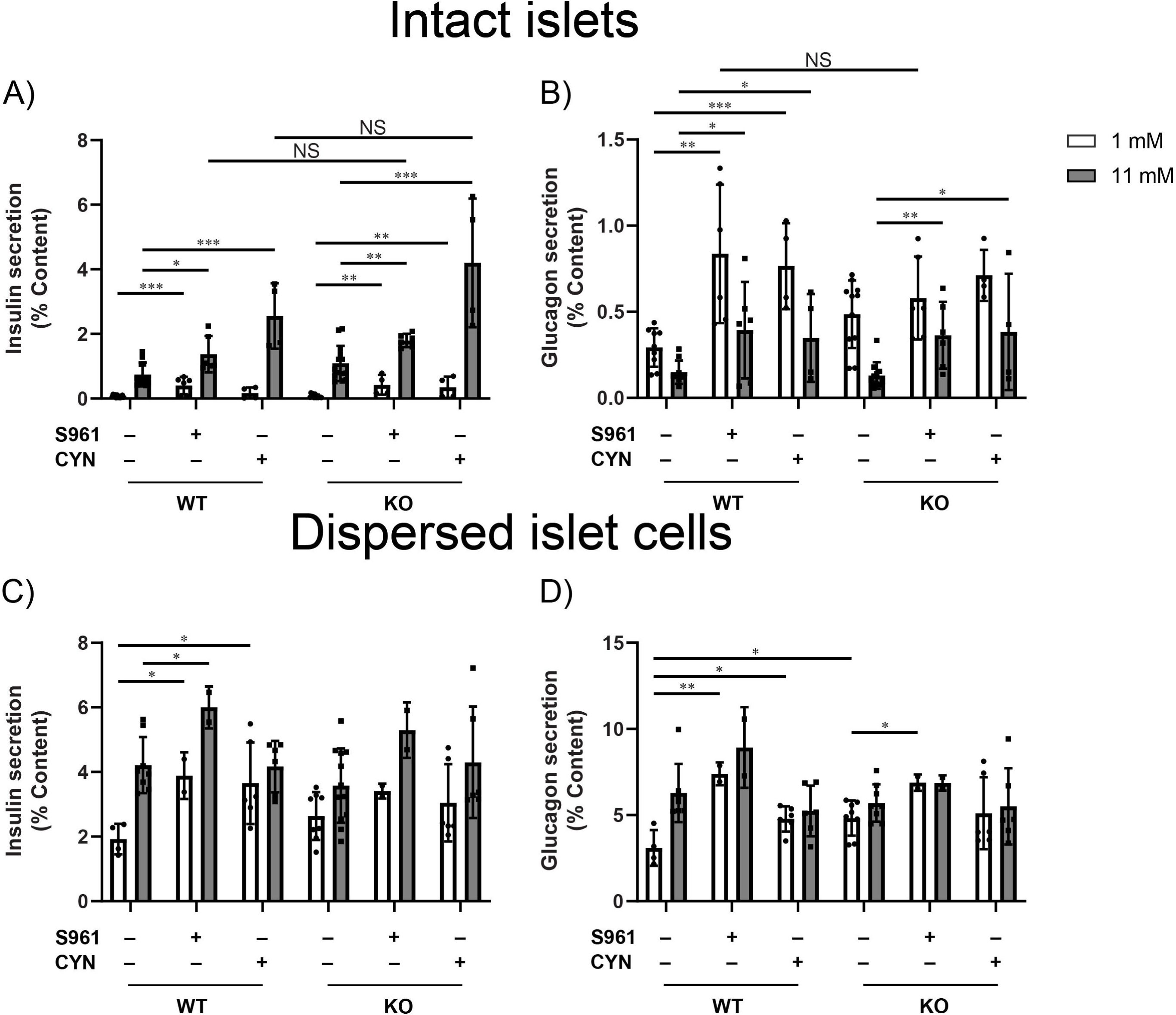
Effects of antagonism of insulin receptor (IR) and type 2 somatostatin receptor (SSTR2) are similar for insulin and glucagon secretion from Cplx 2 KO intact and dispersed mouse islets. Insulin (A) and glucagon (B) secretion from WT and Cplx 2 KO (KO) intact mouse islets at low (1 mM, white) and high (11 mM, gray) glucose concentrations in the presence or absence of 2 µM S961 (IR antagonist) or 100 nM CYN 154806 (CYN, SSTR2 antagonist). Insulin (C) and glucagon (D) secretion from WT and KO dispersed cells at low (1 mM, white) and high (11 mM, gray) glucose concentrations in the presence or absence of 2 µM S961 or 100 nM CYN. Data presented as mean ± SD (n > 2 experiments). Statistical significance between conditions was determined using unpaired *t* test ***P < 0.001; **P < 0.01; *P < 0.05.

### Cplx 2 regulates paracrine inhibition of insulin secretion by somatostatin through β-cell Ca^2+^ activity

Since Cplx 2 has no direct effect on IR and SSTR2 receptor activity, we investigated the downstream pathways that regulate Ca^2+^ influx. We measured Ca^2+^ activity in β- and α-cells of WT and Cplx 2 KO islets in the absence and presence of 1 μM SST at low (1 mM) and high (11 mM) glucose concentrations. To investigate the effects of SST on β-cell Ca^2+^ dynamics, we loaded islets with the Ca^2+^ indicator Fluo-4 AM and identified β-cells based on the glucose response of their Ca^2+^ activity. β-cell Ca^2+^ intensity traces reveal that SST inhibition of β-cell Ca^2+^ activity is absent in Cplx 2 KO islets as compared to the known inhibition of Ca^2+^ dynamics by SST in WT islets (Fig. 5A and B). The percentage change in the normalized Fluo-4 intensity upon 1 µM SST treatment at 11 mM glucose is 59.1 ± 22.7% in WT islets but only 13.5 ± 26.8% in Cplx 2 KO islets (Fig. 5C). Overall, this supports the idea that Cplx 2 is involved in the pathway of paracrine inhibition of insulin secretion by SST downstream of receptor activation, through β-cell Ca^2+^ channel activity.

**Figure 5.**
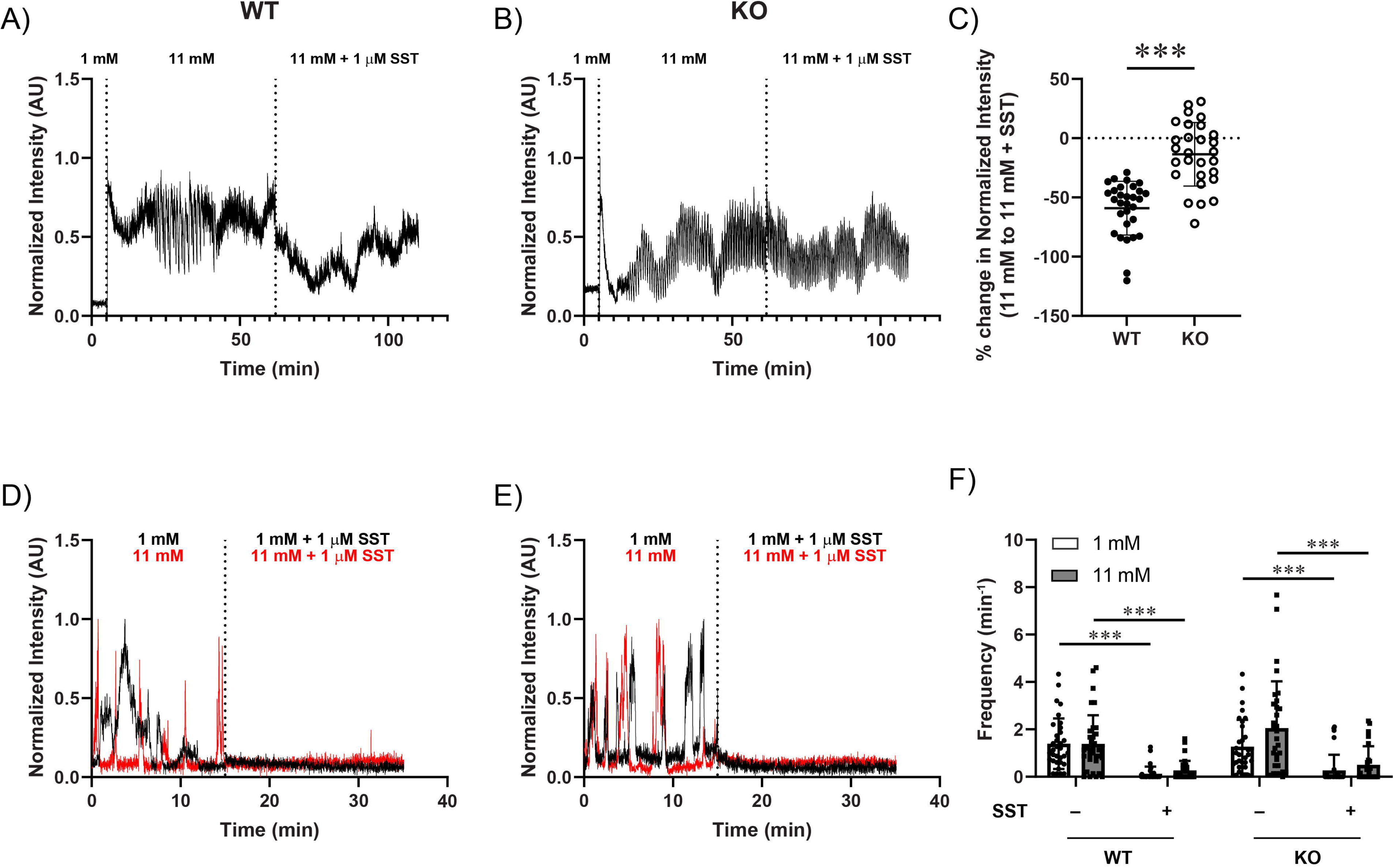
Cplx 2 mediates paracrine regulation of β-cell Ca^2+^ activity by somatostatin. Representative intensity traces of Fluo-4 Ca^2+^ indicator dye in β-cells of WT (A) and KO (B) intact mouse islets with the respective treatment conditions indicated by dashed lines. (C) Percentage change in Fluo-4 intensity averaged over traces acquired at 11 mM glucose concentration in the presence and absence of 1 μM SST for β-cells of WT and KO islets (n > 2 islets, 29 cells). Representative intensity traces of GCaMP6f fluorescence in α-cells of WT (D) and KO (E) islets with treatment conditions indicated by dashed lines. (F) Treatment with 1 μM SST inhibits α-cell Ca^2+^ activity in both WT and KO islets under low (1 mM, white) and high (11 mM, gray) glucose levels (n > 2 islets, 31 cells). Data presented as mean ± SD. Statistical significance between conditions was determined using unpaired *t* test ***P < 0.001; **P < 0.01; *P < 0.05.

The effect of SST treatment on α-cell Ca^2+^ dynamics was also measured in intact WT and Cplx 2 KO islets using α-cell specific GCaMP6f Ca^2+^ biosensor mice (23, 24). Normalized α-cell GCaMP6f intensity traces for WT (Fig. 5D) and Cplx 2 KO (Fig. 5E) islets at low (1 mM, black) or high glucose levels (11 mM, red) before and after 1 µM SST treatment show similar decreases in α-cell Ca^2+^ activity for both WT and Cplx 2 KO islets. The α-cell Ca^2+^ spiking frequency (number of Ca^2+^ peaks per time) also exhibit similar reductions upon treatment with SST for both WT and Cplx 2 KO islets (Fig. 5F). These results are consistent with the glucagon secretion data on intact WT and Cplx 2 KO islets treated with SST (Fig. 3B), where Cplx 2 does not play a role in regulating paracrine inhibition of glucagon secretion by SST through α-cell Ca^2+^ activity.

### Cplx 2 clamps insulin vesicle fusion in β-cells of intact mouse islets through mechanisms involving paracrine inhibition by somatostatin

To assess the role of Cplx 2 in insulin vesicle fusion and exocytosis, we transduced intact WT and Cplx 2 KO mouse islets with the exocytosis biosensor, VAMP2-superecliptic pHluorin (SEP) adenovirus (25, 26). Total internal reflection fluorescence (TIRF) microscopy imaging was performed on WT and Cplx 2 KO islets adhered onto glass bottom dishes with rhLaminin-521 (Fig. 6A and B). β-cells from WT and Cplx 2 KO islets are identified as cells with increased fusion events upon changing from 1 mM to 11 mM glucose, while cells that do not have such a response are not considered. We observe SST inhibition of insulin vesicle fusion events only in β-cells of WT islets and not in β-cells of Cplx 2 KO islets (Fig. 6B). The average half-life (t_1/2_) of vesicle fusion events is longer in β-cells of Cplx 2 KO islets than those of WT islets under all conditions (Fig. 6C). This suggests that loss of Cplx 2 causes irregular vesicle fusion dynamics that may be correlated with the loss of SST inhibition of insulin release.

**Figure 6.**
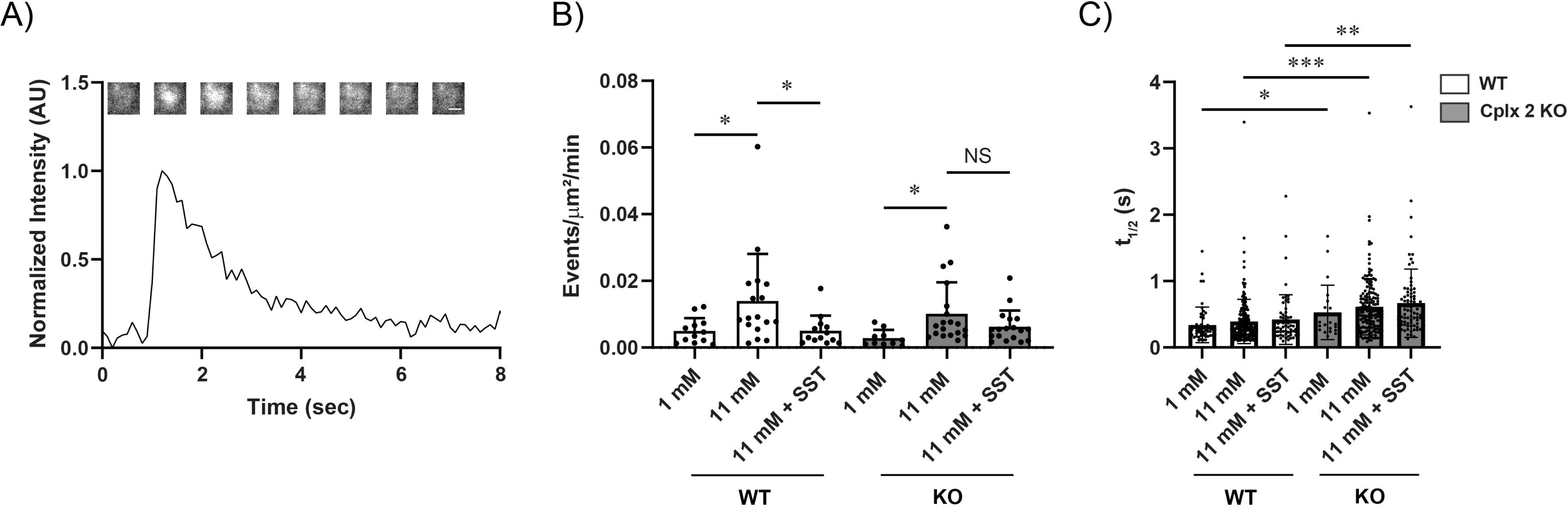
Loss of Cplx 2 dysregulates paracrine inhibition of insulin vesicle fusion by somatostatin. (A) A representative TIRF time trace of an isolated granule fusion event in a VAMP2-superecliptic pHluorin (SEP) adenovirus transduced Cplx 2 KO islet. Scale bar is 1 μm. (B) Insulin granule fusion events in WT and KO VAMP2-SEP islets at low (1 mM) and high glucose (11 mM) concentrations with and without 1 μM SST (n > 2 islets, 11 cells). (C) Half-life (t_1/2_) of fusion events from (B) for WT and KO islets (n > 21 fusion events).

## Discussion

The role of Cplxs have been studied and characterized in neurotransmitter synaptic vesicle fusion and release processes within neurons and neuronal cells (7, 20, 27–29). However, little is known about how Cplxs regulate hormone secretion from pancreatic islet β-, α- and δ-cells. One study explored the role of Cplx 1 in glucose-stimulated insulin secretion with overexpression and siRNA knockdown (21). The authors showed that glucose-stimulated insulin secretion was reduced for both overexpression and knockdown of Cplx 1, which is consistent with it acting as both a negative and positive regulator of insulin secretion. To better understand the role of Cplxs in pancreatic islet cell secretions, we examined the pancreatic islet expression of Cplx 2, the next most highly expressed Cplx in the pancreas after Cplx 1 (9), in β-, α- and δ-cells and explored its function in regulating insulin, glucagon and somatostatin release. Cplx 2 is expressed in pancreatic islet β-, α- and δ-cells as shown by RNA-FISH and immunofluorescence imaging experiments (Fig. 1 and Fig. S1). Loss of Cplx 2 by global knockout results in elevated levels of glucagon and somatostatin secretion from intact islets compared to WT islets (Fig. 2B and C).

These data show that Cplx 2 plays an inhibitory role in clamping α-cell glucagon secretion at low glucose level and δ-cell somatostatin secretion at both low and high glucose levels. In contrast, loss of Cplx 2 does not affect insulin secretion from intact Cplx 2 KO islets (Fig. 2A), likely due to compensatory effects from Cplx 1 which is highly expressed in pancreatic β-cells (21). Such compensatory effect is supported by neuronal studies of excitatory postsynaptic currents (EPSCs) in Cplx 2 single knockout and Cplx 1/2 double knockout mice where peak neurotransmitter release rates of Cplx 1/2 double KO is three times lower than those in Cplx 2 KO cells, and neurotransmitter release from Cplx 1/2 double KO cells is less sensitive to Ca^2+^ than Cplx 2 single KO cells (17). Cplx 2 deficiency in intact mouse islets does not alter the glucose response of insulin, glucagon and somatostatin secretions, suggesting that Cplx 2 plays a secondary role in maintaining proper glucose homeostasis (Fig. 2A – C).

Intercellular communication in islets plays a pivotal role in proper regulation of glucose homeostasis. Dissociation of intact islets into dispersed islet cells changes the functional glucose response of the islet cells (1). This includes a smaller glucose-stimulated insulin secretion response from dispersed β-cells than whole islets (30), a loss of glucose-induced coordinated β-cell Ca^2+^ oscillations and normal two-phase insulin secretion profile in dispersed β-cells (2, 31–33), and reversal of the glucose response of glucagon secretion where glucagon secretion increases with glucose, opposite that of glucose inhibition of glucagon secretion in intact islets and *in vivo* (34–36). Thus, we examined the function of Cplx 2 in dispersed WT and Cplx 2 KO islet cells, which disrupts the normal intercellular communication through paracrine and juxtacrine signaling. We observe slightly elevated glucagon secretion from dispersed Cplx 2 KO cells at low glucose concentrations, but no differences in insulin and somatostatin secretion between dispersed WT and Cplx 2 KO islet cells (Fig. 2D – F). These findings were further substantiated by the lack of difference in insulin and glucagon secretion between dispersed WT and Cplx 2 KO islet cells following treatment with SST or receptor antagonism (Fig. 3C and D and Fig. 4C and D). Taken together, these data provide evidence that the main role of Cplx 2 in the regulation of islet cell secretion requires intercellular communication through paracrine and juxtacrine interactions.

Since we showed that Cplx 2 plays a clamping role in glucagon and somatostatin secretion from intact islets, we examined the correlation of the negative regulatory role of Cplx 2 with paracrine inhibition of insulin and glucagon release by SST. Importantly, the effect of Cplx 2 deficiency on insulin secretion from intact islets was observed only after paracrine inhibition by SST.

Unlike in intact WT islets where SST suppresses insulin secretion at high glucose concentration, intact Cplx 2 KO islets do not respond to treatment with SST (Fig. 3A). This implies that the inhibitory role of Cplx 2 on insulin secretion operates through the paracrine SST inhibition pathway of β-cell insulin secretion. Conversely, glucagon secretion is inhibited by SST treatment at low glucose concentrations for both WT and Cplx 2 KO islets (Fig. 3B), suggesting that Cplx 2 does not play a main role in paracrine inhibition of α-cell glucagon secretion by SST.

Somatostatin, the secretory product of δ-cells, is a major paracrine inhibitor of islet hormone secretion through somatostatin receptors (SSTRs) on β- and α-cells (1, 37). Downstream signaling pathways that suppresses either insulin or glucagon granule exocytosis include: i) the generation of intracellular GTP, causing hyperpolarization of the β or α-cell plasma membrane via the activation of G protein-gated inwardly rectifying K^+^ channels (GIRK), leading to decreased Ca^2+^ influx (38–40); ii) the activation of the Gαi subunit of the type 2 somatostatin receptor (SSTR2) which inhibits adenylyl cyclase and reduces cAMP levels (41, 42); iii) the G_i2_-dependent activation of calcineurin in localized regions where SSTRs associate with voltage-gated L-type Ca^2+^ channels, leading to depriming of glucagon secretory granules close to L-type Ca^2+^ channels (43). Thus, we examined the effect of Cplx 2 at various stages of the paracrine signaling pathway in β- and α-cells, starting with the membrane receptors involved in paracrine inhibition of insulin and glucagon secretions. We observe no significant differences between WT and Cplx 2 KO islets in terms of insulin and glucagon secretion following antagonism of the IR with S961 or SSTR2 with CYN 154806 (Fig. 4A and B). These data indicate that the effect of Cplx 2 likely occurs downstream of membrane receptor activation.

One signaling pathway underlying SST inhibition of insulin and glucagon secretion is the inactivation of voltage-gated Ca^2+^ channels and the consequent decrease in intracellular Ca^2+^ influx. We examined the Ca^2+^ activities of both β- and α-cells in intact WT and Cplx 2 KO islets before and after treatment with SST to test a potential role of Cplx 2 in regulating Ca^2+^ channel activity. For β-cell Ca^2+^ activity, WT and Cplx 2 KO islets were loaded with Ca^2+^ indicator dye Fluo-4 AM and time-series imaging was conducted. β-cells were identified as islet cells with negligible Ca^2+^ dynamics at low glucose level that became active upon a switch to 11 mM glucose. The time series data reveal that SST inhibition of β-cell Ca^2+^ activity is lost in Cplx 2 KO islets (Fig. 5A and B), which is consistent with the lack of SST treatment effect on insulin secretion in Cplx 2 KO islets (Fig. 5C and Fig. 3A). These data confirm that Cplx 2 mediates SST inhibition of insulin secretion by regulating β-cell Ca^2+^ activity. In contrast, α-cell Ca^2+^ activity (measured as frequency of bursts of Ca^2+^ influx events) is suppressed by SST treatment in both WT and Cplx 2 KO islets regardless of glucose concentration (Fig. 5D – F). These data match the glucagon secretion results of SST-treated WT and Cplx 2 KO islets (Fig. 3B), indicating that Cplx 2 is not involved in the regulation of paracrine inhibition of α-cell Ca^2+^ activity and glucagon secretion by SST.

Since Cplx 2 is directly involved in the vesicle fusion and release process through its binding to the prefusion SNARE protein complex (5, 6, 10), we measured insulin vesicle fusion events using VAMP2-SEP. This biosensor is quenched in acidic vesicles (pH < 6) and exhibits a burst of fluorescence during vesicle fusion followed by subsequent fluorescence decay upon vesicle release (Fig. 6A). We used TIRF imaging to visualize vesicle fusion events in WT and Cplx 2 KO islets attached to rhLaminin-521-coated glass bottom dishes. Consistent with the insulin secretion and β-cell Ca^2+^ activity measurements, SST treatment does not inhibit insulin vesicle fusion events in Cplx 2 KO islets (Fig. 6B). Notably, the half-life of β-cell vesicle fusion events in Cplx 2 KO islets is significantly longer than that of WT islets under all conditions (Fig. 6C).

This is consistent with a loss of Cplx 2 clamping effects on the prefusion SNARE protein complex, which would be expected to cause aberrant vesicle fusion pore opening and release kinetics. Taken together, these data confirm the role Cplx 2 plays in paracrine regulation of insulin secretion in intact islets through β-cell Ca^2+^ activity.

Our study demonstrates that Cplx 2 plays a direct inhibitory role in glucagon and somatostatin secretion and indirectly regulates insulin secretion through SST paracrine effects that modulate β-cell Ca^2+^ activity and vesicle release kinetics in intact mouse islets. However, we acknowledge that possible compensatory effects from Cplx 1 could play a role, especially in β-cells (21). Complications from Cplx 1 activity, though, should not alter the inhibitory role of Cplx 2 established here since compensatory effects from Cplx 1 would be expected to mask the effects of Cplx 2 deletion and reduce the magnitude of any effects. Further studies with Cplx 1 or Cplx 2 single KO and Cplx 1/2 double KO mouse lines (17) targeted specifically to islet cell types (24, 44) could be used to study the complimentary and independent effects of Cplx 1 and Cplx 2. It should be noted that global double KO of Cplx 1/2 is lethal in mice embryos (17). In addition to Cplx 2 interactions with β-cell Ca^2+^ activity, it could also interact with another β-cell second messenger, cAMP, to regulate insulin secretion. It was reported that Cplx interacts with cAMP through protein kinase A (PKA) activation, which subsequently phosphorylates Cplx and augments spontaneous neurotransmitter release in *Drosophila* (28). This pathway is similar to what has been shown for SST treatment of pancreatic α-cells (41), although we did not find a role for Cplx 2 consistent with those findings. Further studies on the interaction between Cplx 2 and cAMP could complete the mechanistic pathway of Cplx 2 in regulating islet cell secretions, especially for the β-cell. Lastly, several papers highlight the role of different proteins involved in the SNARE protein complex assembly and disassembly in β-cells, such as Munc-18, Syntaxin 1, synaptotagmin-7, and others (45–48). A complete understanding of Cplx 2 in regulating the secretion of insulin, glucagon and somatostatin from pancreatic islets will require insights into its dynamic interactions with these other proteins that are directly (t-SNARE: synaptosome associated protein 25 (SNAP-25) and syntaxin-1 found on the plasma membrane and v-SNARE: vesicle associated membrane protein 2 (VAMP2) and synaptotagmin localized on the vesicle membrane) and indirectly (Munc-18, Stxbp) involved in the assembly and disassembly of the SNARE protein complex.

## Methods

### Islet Isolation and Culture

All animal experiments were performed under approval of the Washington University Institutional Animal Care and Use Committee (IACUC). Briefly, murine islets were isolated using 0.075% collagenase digestion at 34□ post pancreatectomy, picked manually under a stereomicroscope, cultured in islet media (RPMI 1640 with 10% FBS, 11 mM Glucose, and penicillin-streptomycin) at 37□ /5% CO_2_ overnight before use (35). Homozygous Cplx 2 knockout (KO) mutant mice sperm were received as a kind gift from Dr. Reim in the Max Planck Institute for Multidisciplinary Sciences and in-vitro fertilization (IVF) was performed on wild-type (WT) female mice of C57BL/6 background strain. After several generations of breeding between heterozygous Cplx 2 mutant mice, Cplx 2 KO mice were eventually derived and used for experiments. Islets with α-cells expressing GCaMP6f were obtained by crossing mice expressing a Cre-dependent GCaMP6f (JAX # 028865) with mice expressing a Glucagon-iCre (JAX # 030663). Further crossing between GCG-iCre-GCaMP6f mice with homozygous Cplx 2 KO mutant mice generated GCG-iCre-GCaMP6f Cplx 2 KO mice.

### Preparation of dispersed islet cells

Isolated islets were washed with Hanks’ balanced salt solution (without Ca^2+^ or Mg^2+^, pH 7.4, Gibco) then dissociated in Accutase (Innovative Cell Technologies) for 15 minutes at 37□ with intermittent trituration. Dissociated cells were then resuspended in islet culture media for subsequent use. For secretion assays, dispersed islet cells were split into microcentrifuge tubes for equilibration and the respective treatment conditions.

### Secretion Assays

Islets (8 islets per condition for insulin or glucagon secretion assay; 30 islets per condition for somatostatin secretion assay) or dispersed islet cells (15 islets per condition for insulin or glucagon secretion assay; 30 islets per condition for somatostatin secretion assay) were first equilibrated in KRBH buffer (128.8 mM NaCl, 4.8 mM KCl, 1.2 mM KH_2_PO_4_, 1.2 mM MgSO_4_·7H_2_O, 2.5 mM CaCl_2_, 20 mM HEPES, and 5 mM NaHCO_3_, pH 7.4) with 0.1% BSA and 2.8 mM glucose for 45 minutes at 37° C. Thereafter, islets or dispersed islet cells were incubated in 100 µl (insulin and glucagon secretion assays) or 150 µl (somatostatin secretion assay) or KRBH buffer at low (1 mM) or high (11 mM) glucose levels with either 1 µM somatostatin (SST, Millipore Sigma), 2 µM S961 (Phoenix Pharmaceuticals Inc.) or 100 nM CYN 154806 (Fisher Scientific) treatment in 1.5 ml microcentrifuge tubes for 1 hour at 37°C. The supernatant was then collected for secretion measurement. Islets were then transferred into microcentrifuge tubes containing 100 µl acidified ethanol solution (1.5% HCl in 70% ethanol), or in the case of dispersed islet cells, resuspended in 100 µl acidified ethanol solution, for overnight cell lysis and hormone content extraction at -20°C. Insulin and glucagon concentrations were measured by Lumit Insulin and Glucagon Immunoassay kits (Promega), while somatostatin concentration was measured with ELISA (Phoenix Pharmaceuticals Inc.). Data are presented as percent total hormone content as a normalization for sample differences.

### Fixed pancreatic islet section immunohistochemistry assay

WT and Cplx 2 KO mice pancreases were fixed in 10% neutral buffered formalin solution overnight at room temperature and sent to the Anatomic and Molecular Pathology Core labs in Washington University in St. Louis for paraffin-embedding and sectioning to obtain paraffin-embedded pancreatic sections. The paraffin-embedded pancreas sections were deparaffinized with sequential incubation twice in xylene (10 minutes), once in 100% ethanol (5 minutes), once in 95% ethanol (5 minutes), once in 70% ethanol (5 minutes) and once in deionized water (2 minutes). After deparaffinization, antigen retrieval was performed by boiling the pancreas sections with Diva Decloaker buffer (Biocare) for 20 minutes. The pancreas sections were then cooled and permeabilized with 0.1% Triton X-100 in PBS-T (phosphate buffered saline (PBS) containing 0.05% Tween 20) for 30 minutes at room temperature and blocked in 2% BSA in PBS-T blocking buffer for 1 hour. Pancreas sections were then incubated with primary antibodies diluted in blocking buffer against either insulin (2D11-H5; 1:100; Santa Cruz), glucagon (MAB1249; 1:100; R&D Systems) or somatostatin (MAB2358; 1:50; R&D Systems) along with Cplx 2 (18149-1-AP; 1:50; Proteintech) overnight at 4°C. This is followed by incubation with the corresponding secondary antibodies (1:1000 Alexa Fluor 594 goat anti-mouse for glucagon and insulin, 1: 1000 Alexa Fluor 594 goat anti-rat for somatostatin; and 1:1000 Alexa Fluor 488 goat anti-rabbit for Cplx 2, ThermoFisher Scientific) for 1.5 hour at room temperature. Stained pancreas section slides were then washed in PBS, mounted in ProLong Glass Antifade Mountant (Invitrogen) and imaged on the Zeiss LSM 880 confocal microscope with a Plan-Apochromat 40×, 1.3 NA oil objective lens, a multi-band beamsplitter for 488/561/633 laser excitations, sequentially excited by 488 nm and 561 nm excitation lines and sequentially detected at emission ranges of 500 – 552 nm (Alexa Fluor 488) and 596 – 686 nm (Alexa Fluor 594) using spectral GaAsP detectors.

### RNA Fluorescence in situ hybridization (FISH)

RNA-FISH experiments were conducted in fixed WT mouse islets according to the manufacturer’s instructions in the RNAScope Multiplex Fluorescent v2 Assay combined with immunofluorescence workflow (#323100, ACDBio). Isolated WT islets were cultured overnight on glass bottom dishes coated with 1.5 µg/cm^2^ rhLaminin-521 (Gibco). Islets were then fixed with 2% paraformaldehyde for 20 minutes, permeabilized in 0.1% Tween 20 in PBS for 1 hour and blocked in blocking buffer (2% BSA in PBS-T) for 1 hour at room temperature. Fixed islets were subsequently hybridized with custom-designed *Mus musculus* Cplx 2 (Mm-Cplx 2-C3) probe and developed with Opal-520 (FP1487001KT; 1:1000; Akoya Biosciences). Fixed islets were then immunostained with primary antibodies against either insulin (2D11-H5; 1:100; Santa Cruz), glucagon (MAB1249; 1:100; R&D Systems) or somatostatin (MAB2358; 1:50; R&D Systems) and their corresponding secondary antibodies, as detailed above. DAPI staining was conducted according to the RNAScope Multiplex Fluorescent v2 Assay combined with immunofluorescence workflow for identification of the cell nuclei. Fixed islets were then imaged on the Zeiss LSM 880 confocal microscope with a Plan-Apochromat 63×, 1.4 NA oil objective lens, a multi-band beamsplitter for 405 and 488/561/633 laser excitations, sequentially excited by 405nm, 488 nm and 561 nm excitation lines and sequentially detected at emission ranges of 410 – 476 nm (DAPI), 500 – 552 nm (Opal 520) and 596 – 686 nm (Alexa Fluor 594) using spectral GaAsP detectors.

### Live islet calcium imaging

For β-cell calcium imaging, WT and Cplx 2 KO islets were first cultured in islet media for 2 days on glass bottom dishes coated with 1.5 µg/cm^2^ rhLaminin-521. Islets were then incubated with 4 µM Fluo-4 AM for 45 minutes at 37°C/5% CO_2_ in 2 mM glucose-containing KRBH buffer (pH 7.4) with 0.1% BSA. Next, islets were pre-incubated in 1 mM glucose-containing KRBH buffer (pH 7.4) with 0.1% BSA for 5 minutes in a temperature and CO_2_ controlled stage heater at 37°C/5% CO_2_ prior to imaging. Time-lapse confocal imaging of the Fluo-4 AM loaded islets was conducted on the Zeiss LSM 880 confocal microscope with a Plan-Apochromat 63×, 1.4 NA oil objective lens, a multi-band beamsplitter for 488/561/633 laser excitations, with 488 nm laser excitation and detected at an emission range of 500 – 597 nm, at a frame rate of 1.6 Hz.

For α-cell calcium imaging, GCG-iCre-GCaMP6f WT and GCGiCre-GCaMP6f Cplx 2 KO mice islets were cultured in islet media for 2 days on glass bottom dishes coated with 1.5 µg/cm^2^ rhLaminin-521. Islets were equilibrated in 2.8 mM glucose-containing KRBH buffer (pH 7.4) with 0.1% BSA for 15 minutes at 37°C/5% CO_2_. Thereafter, islets were pre-incubated in either 1 mM or 11 mM glucose-containing KRBH buffer (pH 7.4) with 0.1% BSA for 30 minutes on the temperature and CO_2_ controlled stage heater at 37°C/5% CO_2_ prior to imaging. Time-lapse confocal imaging was then conducted for α-cell GCaMP6f-labeled islets on the Zeiss LSM 880 confocal microscope with the same settings that were used for β-cell calcium imaging.

### TIRF imaging of insulin vesicle fusion in intact islets

WT and Cplx 2 islets were cultured and transduced with VAMP2-superecliptic pHluorin (SEP) adenovirus (Vector Biolabs) in glass bottom dishes coated with 1.5 µg/cm^2^ rhLaminin-521 for 2 days. At least 1 hour prior to imaging, the virus-containing islet media was replaced with virus-free islet media in the glass bottom dishes. Next, islets were equilibrated in 2.8 mM glucose-containing KRBH buffer (pH 7.4) with 0.1% BSA for 15 minutes at 37°C/5% CO_2_. Islets were pre-incubated in 1 mM glucose-containing KRBH buffer (pH 7.4) with 0.1% BSA for 15 minutes on a temperature and CO_2_ controlled stage heater at 37°C/5% CO_2_ prior to imaging. Time-lapse TIRF imaging was performed on a Nikon TIRF microscope system with an Apo TIRF 100×, 1.49 NA oil objective lens, laser excitation at 488 nm, at an exposure time of 100 ms on the ORCA-Fusion BT Digital sCMOS camera (Hamamatsu). VAMP2-SEP fusion events were manually identified and analyzed based on the rapid fluorescence burst, which is immediately followed by an exponential fluorescence decay of the VAMP2-SEP labeled vesicles. Half-life (t_1/2_) of the fusion events were extracted by fitting the fluorescence fusion decay curve to a single exponential decay function.

### Data analysis and Statistics

Data were analyzed in Microsoft Excel, ImageJ, and GraphPad Prism and Matlab. α-cell Ca^2+^ imaging data was analyzed by counting the number of peaks using a peakfinding script that is available in Matlab and dividing with the imaging time to obtain the average Ca^2+^ frequency. Data are reported as mean values ± SD. Statistical significance was assessed as described in the figure captions.

### Data and resource availability

The data set generated during or analyzed during the current study is available from the corresponding author on reasonable request.

## Acknowledgements

The authors thank Dr. Kerstin Reim and Prof. Dr. Nils Brose (Max Planck Institute for Multidisciplinary Sciences) for providing the homozygous Cplx 2 knockout (KO) mutant mice line. The authors thank the Mouse Genetics Core and the Anatomic and Molecular Pathology Core labs in Washington University in St. Louis for their services in IVF and paraffin-embedding and sectioning, respectively.

## Author contributions

XWN, MRD and DWP designed and planned the research. XWN, MRD, CK and JL performed experiments, where XWN and MRD conducted analysis. XWN and DWP wrote the first draft of the manuscript, which was edited and approved by all authors. DWP is the guarantor of this work and, as such, had full access to all the data in the study and takes responsibility for the integrity of the data and the accuracy of the data analysis.

## Figures

**Figure S1.** (A) Quantification of Cplx 2 RNA copy per cell in insulin (β-cell), glucagon (α-cell) and somatostatin (δ-cell) immunostained fixed WT mouse islet (n > 2 cells). (B) Mean fluorescence intensities of Cplx 2 antibody co-immunostained with either insulin (β-cell), glucagon (α-cell) or somatostatin (δ-cell) antibodies in WT and Cplx 2 KO mouse pancreatic islet sections (n > 4 islets).

