## Supplemental Figure 1 for "Role of Complexin 2 in the regulation of hormone secretion from the islet of Langerhans"

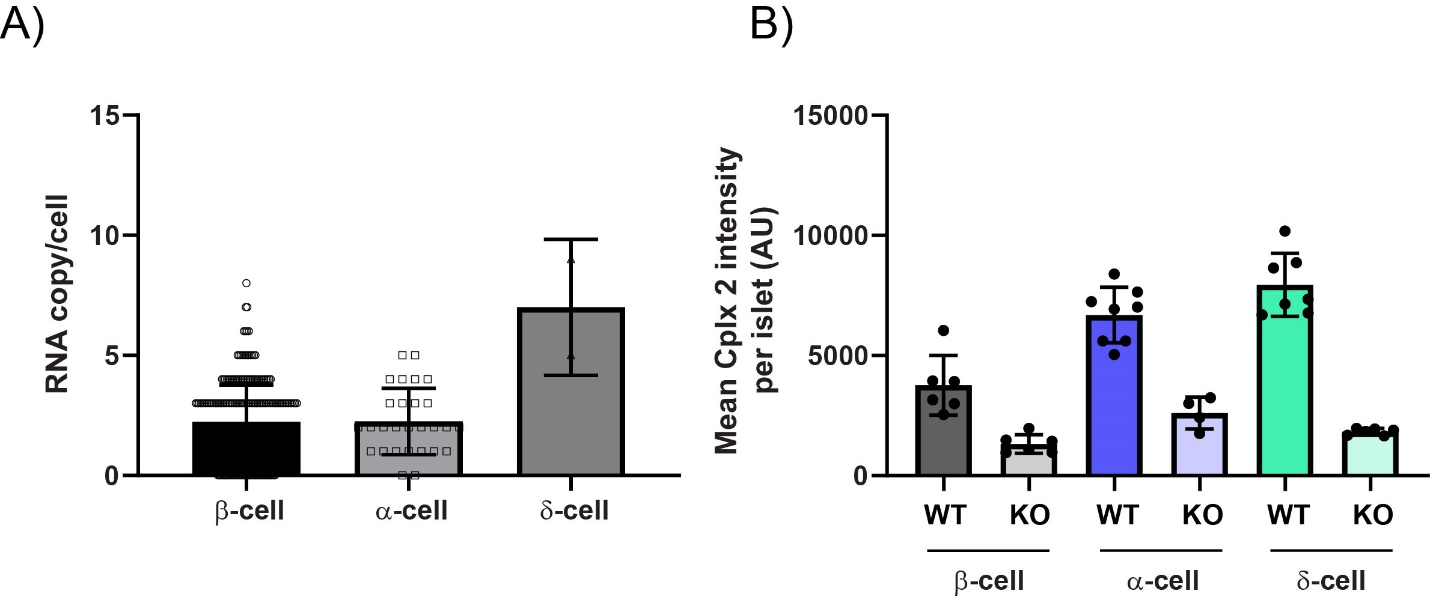

**Figure S1.**  (A) Quantification of Cplx 2 RNA copy per cell in insulin (β-cell), glucagon (α-cell) and somatostatin (δ-cell) immunostained fixed WT mouse islet (n > 2 cells). (B) Mean fluorescence intensities of Cplx 2 antibody co-immunostained with either insulin (β-cell), glucagon (α-cell) or somatostatin (δ-cell) antibodies in WT and Cplx 2 KO mouse pancreatic islet sections (n > 4 islets).
